# Force regulates mitotic entry timing during Drosophila gastrulation

**DOI:** 10.64898/2026.11.12.766297

**Authors:** Mingmar Sherpa, Thomas Evans, Adam C. Martin

## Abstract

Cell division and cell shape changes drive tissue morphogenesis. Cell division is tightly regulated by biochemical factors that influence the cell cycle. Past work in our lab showed how contractility of mesoderm cells influences cell division orientation during *Drosophila* gastrulation. Whether contractile forces that drive morphogenetic folding influence the timing of coincident mitotic divisions is unknown. During *Drosophila* gastrulation, cell division is regulated by mitotic entry of cells in stereotypic regions across the embryo, termed mitotic domains. Here, we examine the role of morphogenesis in the timing of mitotic entry in the mitotic domains. To do this, we disrupted adherens junctions to globally disrupt force transmission across the embryo. In addition, we examined the timing of mitotic entry in mutants that affect mesoderm invagination. Our data suggests a role for mesoderm invagination in mitotic domain 1 timing.

## Introduction

Cell growth and division are essential components of morphogenesis. In its simplest form, cell growth and proliferation lead to tissue expansion (Burda *et al*., 2023). Oriented cell growth or division patterns can lead to tissue extension, such as those that occur during the axis of vertebrates (Concha and Adams, 1998). Differential growth between two attached tissues can lead to more complicated deformations such as folds and loops (Savin *et al*., 2011; Shyer *et al*., 2013; Yin *et al*., 2025). Thus, regulation of cell division patterns in the embryo plays a critical role in the resulting shapes of tissues.

The cell cycle is regulated by both chemical and mechanical signals during morphogenesis. Chemical signals, such as morphogens, are critical for promoting spatial and temporal patterns of cell growth (Wartlick *et al*., 2011). In addition to these signals, mechanical cues can modulate patterns of growth (Shraiman, 2005). For example, tissue stiffness, matrix stiffness, and the spread area of a cell regulate cell cycle progression in vertebrates. And the *Drosophila* wing disc, tensile stresses in the tissue feedback to control cell cycle patterns via the Hippo signaling pathway (Alégot *et al*., 2019; Tokamov *et al*., 2023). Hippo, signaling also responds to mechanics in higher metazoans (Piccolo *et al*., 2014). Other control points in the cell cycle, including the G2 to M transition, have also been shown to respond to force (Gudipaty *et al*., 2017). Whether and how these forms of mechanical control coordinate morphogenetic events in an embryo is poorly understood.

In the *Drosophila* embryo, the cell cycle is precisely controlled in space and time.

After fertilization, nuclei in the syncytial embryo undergo 13 rounds of synchronous division followed by a cell cycle pause in G2 during which the blastoderm forms (Foe, 1989). Subsequently, groups of contiguous cells, termed mitotic domains, enter mitosis in a stereotypic spatial and temporal pattern. Mitotic domains 1 and 5 are the 1^st^ and 5^th^ group to exit G2 and occur in the dorsal head when tissue folding events are happening in different regions of the embryo. It has been shown that the timing and location of these mitotic domains depend on the gene expression patterns of *string*, a *Drosophila* Cdc25 homologue (Edgar and O’Farrell, 1989, 1990). Indeed, it has been shown that the precise timing of mitotic entry is linked to String expression level, with divisions occurring as a wave across the mitotic domain (Di Talia and Wieschaus, 2012). Roles for external cues in the timing of these divisions have not yet been described.

*Drosophila* gastrulation involves an intricate choreography of different cell behaviors and tissue movements (Leptin, 2005; Martin, 2020). Prominent in gastrulation are the tissue folds that occur in precise embryonic positions. The presumptive mesoderm folds inward and is internalized at the ventral midline, which results in ventrally directed tissue movement in the embryo trunk (Butler *et al*., 2009; Lye *et al*., 2015; Rauzi *et al*., 2015; Gustafson *et al*., 2022). Both posterior midgut invagination and trunk cell intercalation lead to posterior cell movement in the trunk and trunk elongation (Butler *et al*., 2009; Collinet *et al*., 2015). Between the head and trunk regions, the cephalic furrow forms at ∼ 30% egg length, which serves as a mechanical buffer to cell movement (Dey *et al*., 2025; Vellutini *et al*., 2025). As a result of these folds, cells are lost from this embryo surface, which is compensated for, in part, by cells rounding upon entering mitosis (Stern *et al*., 2022).

It is likely that mechanical forces play a role in coordinating tissue movements and cell behaviors across the embryo. However, the current model is that these divisions are hard wired to occur through the expression of the *string* gene. Cells in several of the mitotic domains undergo oriented division, which has been shown to depend on forces from surrounding morphogenetic events (da Silva and Vincent, 2007; Wang *et al*., 2017; Scarpa *et al*., 2018; Camuglia *et al*., 2022). Spindle orientation in the embryo depends on a combination of molecular and physical cues, such as cell shape (Chanet *et al*., 2017). Tissue folding was also demonstrated to be coordinated with axis extension. Pulling forces from the mesoderm result in the accumulation of myosin II in the germband (Gustafson *et al*., 2022). This patterned mechanical feedback is required for mesoderm internalization and depends on EGF activity in the germband (Clarke *et al*., 2025).

Here, we used extended time lapse imaging to define the timing of the cycle 14 divisions associated with gastrulation. We found that α-catenin depletion affects the onset of the MD1 and MD5 mitotic entry, possibly by accelerating cellularization. In addition, α-catenin depletion and blocking tissue folding events affected the synchrony of divisions within MD1, but not MD5. We propose that forces transmitted from morphogenetic processes ensure that division timing happen in a coordinated fashion.

## Results

### Morphogenetic events and cell cycle control exhibit robust timing, with respect to each other

To investigate the coordination of ectodermal cell behaviors during *Drosophila* gastrulation, we measured the relative timing of mitotic entry and morphogenetic movements with respect to Anaphase 13. We examined time lapse images of embryos expressing Histone::GFP and measured the timing of the cephalic furrow relative to anaphase 13 and mitotic domains 1 and 5 (MD1 and MD5). We found that the onset of cephalic furrow initiation occurred within a 7 min time window, ∼ 67 minutes following anaphase 13 (Fig. 1A, B, and C). Furthermore, the onset of MD1 rounding occurred with a reproducible time after cephalic furrow initiation (Fig. 1D and 1F). Finally, MD5 initiated a few minutes following MD1 (Fig. 1E). The stereotypic timing of each of these behaviors suggested the presence of mechanisms that coordinate these events.

**Figure 1.**
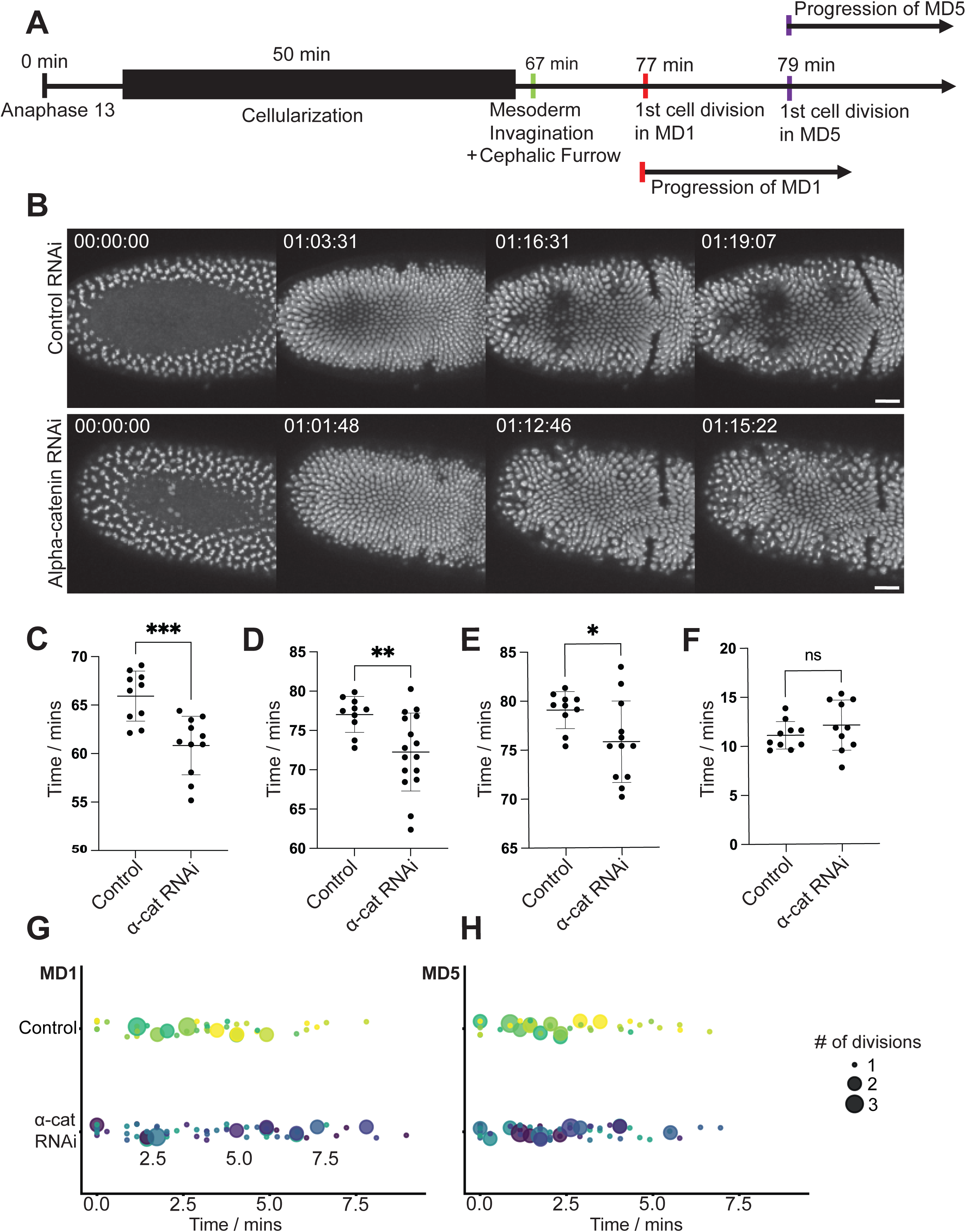
Alpha-Catenin RNAi affects timing of cell division and synchrony of Mitotic Domains. (A) Cephalic furrow, MD1, and MD5 divisions occur with precise timing relative to Anaphase 13 and to each other. (B) Montage of time-lapse images of embryos expressing H2B::GFP. Compared are control knock-down and α-catenin depleted embryos. Scale bar = 25µm. (C) α-catenin depletion accelerated CF initiation. Time of CF initiation relative to Anaphase 13 calculated for 10 control and 11 α-catenin depleted embryos expressing H2B::GFP. Statistical significance was assessed using an unpaired Student’s t-test. (D, E) α-catenin depletion accelerated MD onset. Time of MD1 (D) or MD5 (E) onset relative to Anaphase 13 calculated for control and α-catenin depleted embryos expressing H2B::GFP. (F) MD1 timing relative to CF initiation is unchanged after α-catenin depletion. Time between CF initiation and first MD1 division. (G) Synchrony of MD1 divisions is disrupted by α-catenin depletion in some embryos, whereas (H) MD5 divisions are unaffected. Embryos expressing H2B::GFP were used to calculate the time required for first 10 divisions on the left side of the embryo. In population comparisons, disruption to MD1 division synchrony was not statistically sigificant showing a 30.4% increase in variabilty (p=0.192) and and increase in mean division time of 0.4 minutes (p=0.48) compared to control. Variabilty was measured by comparing the standard deviation of division times within each embryo by genotype conditions. Mean division time was compared using linear-mixed effects models that accounted for multiple divisions being measured within each embryo.

### α-catenin depletion affects the timing of cephalic furrow, MD1, and MD5 initiation and MD1 mitotic entry synchrony

The mechanical linkage between cells could be important for coordinating behavior. Because adherens junctions play a central role in connecting cells, we examined the effects of disrupting adherens junctions on the timing of developmental events. α-catenin depletion lowers the levels of all core adherens junction proteins throughout the *Drosophila* embryo (Denk-Lobnig *et al*., 2021) and has been shown to result in severe cytoskeletal connectivity defects (Yevick *et al*., 2019; Eritano *et al*., 2020). Therefore, we hypothesized that any behaviors whose timing required mechanical coordination across the tissue would be changed upon α-catenin loss.

Prior work showed that α-catenin depletion can disrupt the cephalic furrow (Eritano *et al*., 2020; Popkova *et al*., 2024). Unexpectedly, we found that depleting α-catenin also accelerated cephalic furrow initiation relative to anaphase 13 (Figure 1B and 1C). α-catenin depletion also accelerated the onset of the first cell division in MD1 and MD5 (Figure 1D and 1E). Interestingly, the timing of MD1 with respect to the cephalic furrow initiation was the same as wild type in the α-catenin depletion (Figure 1F). We also measured the timing of mitotic entry within individual domains. Mitotic domains progress as a wave, with the timing of divisions related to String expression (Di Talia and Wieschaus, 2012). Although the population averages were not statistically different, we found that, in some embryos, α-catenin depletion increased the time for cell divisions to occur sequentially within MD1 which we refer to as a disruption in ‘mitotic domain synchrony’ (Figure 1G). We did not observe a change in the propagation of division in MD5. Thus, α-catenin is required for the proper timing of cephalic for initiation and MD1 and MD5 mitotic onset and may affect the synchrony of MD1. We therefore sought to investigate morphogenetic programs that might influence the timing of these events.

### α-catenin depletion changes the duration of cellularization

We next tested whether the advance in cephalic furrow initiation and mitotic domain initiation could be the result of more rapid earlier developmental events. We examined the effect of α-catenin depletion on cellularization. We made brightfield movies of control and α-catenin-RNAi embryos and measured the rate of furrow canal ingression.

α-catenin-RNAi resulted in more rapid furrow canal ingression during fast phase of cellularization (Figure 2A, 2B, and 2B’). Despite the increased rate of cellularization, the depth that the furrow canals reached was indistinguishable from that of control embryos (Figure 2C and 2D). The anterior-posterior position of the cephalic furrow remained unchanged (Figure 2E). Our data suggests that advanced cephalic furrow initiation and mitotic entry in α-catenin depleted embryos is the result of more rapid cellularization. We next investigated whether mesoderm internalization movements could affect the timing of MD1 and MD5 in the dorsal head.

**Figure 2.**
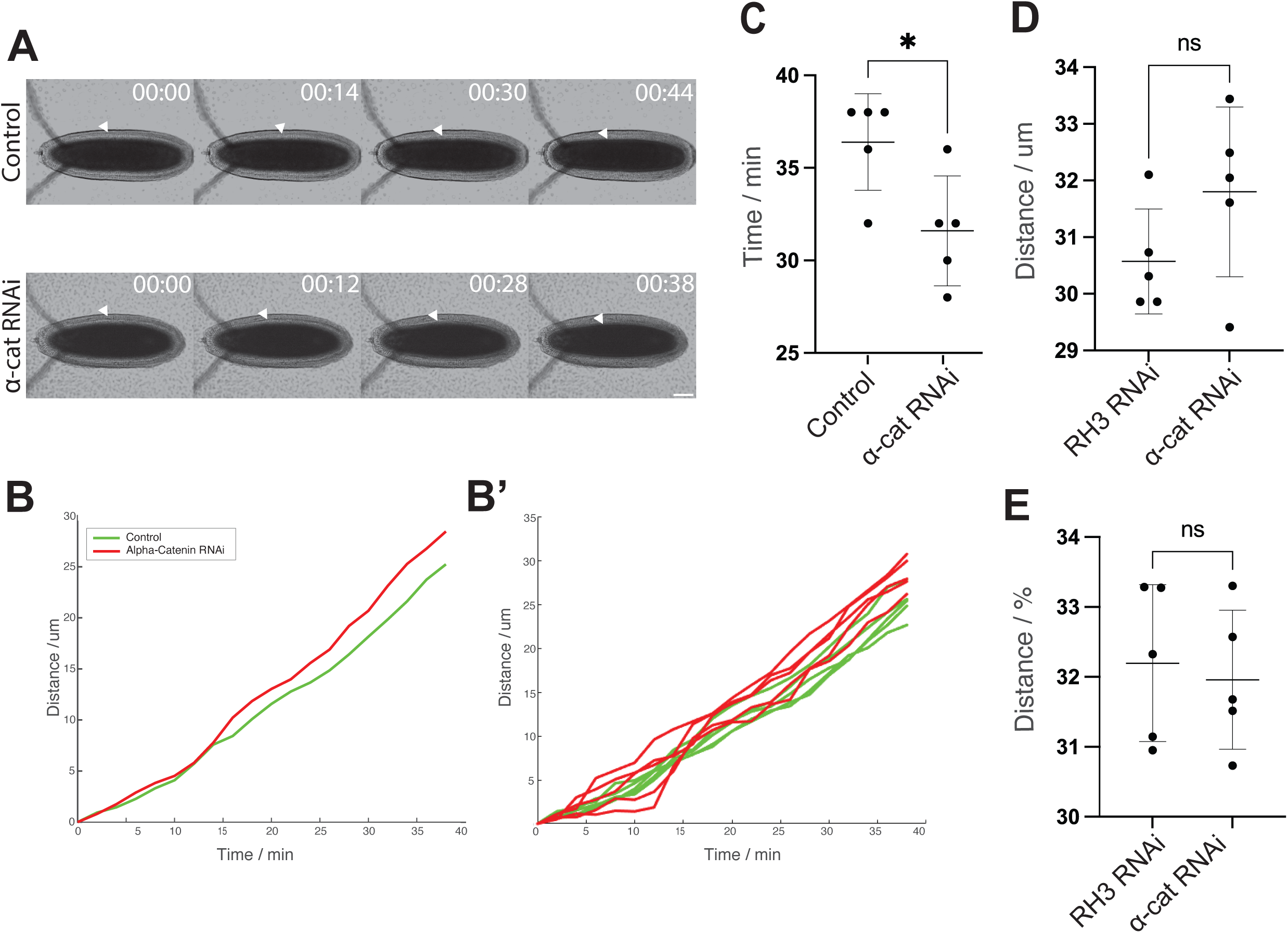
α-catenin depletion affects cellularization rate. (A) α-catenin depletion increases cellularization rate. Brightfield movies tracking cellularization front in control and α-catenin depleted embryos. Scale bar = 25 µm. (B) Plot is the mean furrow canal depth over time measured using brightfield microscopy and aligned by the onset of cellularization. (B’) Shows time traces of individual embryos used to calculate mean cellularization rate in (B). (C) Time for furrow canals to descend 25 µm during fast phase of cellularization. (D) α-catenin depletion does not affect cell height. Cell height calculated from brightfield movies and time of CF initiation. (E) α-catenin depletion does not affect CF position. Anterior-posterior position of CF was measured relative to embryo length. Same statistics as (C). (C-E) Each data point represents one embryo. Five embryos were analyzed per group and statistical significance was assessed using an unpaired Student’s t-test.

### MD1 synchrony is disrupted in *snail* mutant

Mechanical forces produced by coordinated apical constriction drive the formation of the ventral furrow, generating tissue-level tension required for mesoderm invagination. The transcription factor Snail is essential for this morphogenetic movement, as it defines the presumptive mesoderm and regulates the cellular behaviors underlying apical constriction. Importantly, *snail* is expressed in ventral midline cells, not dorsal cells, and in contrast to the *twist* mutant disrupts Myosin II pulses in addition to Myosin II network formation (Martin *et al*., 2009). We used homozygous *snail* mutants to disrupt ventral furrow formation and the mechanical forces associated with mesoderm invagination.

While we initially observed defects using Histone::GFP, we eventually found it useful to select *snail* mutants unambiguously using a linked marker (see Methods) and found that Myosin II imaging made it easier to visually illustrate when cells had undergone division. We then investigated whether the disruption of the ventral furrow influenced the synchrony of mitotic entry in MD1 and MD5. We found that the *snail* mutant disrupted the synchrony of MD1 divisions, increasing the amount of time for the first 10 divisions (Figure 3A and 3B). Like α-catenin depletion the disruption in mitotic synchrony was most apparent for MD1 – there was no significant effect on MD5 division timing (Figure 3A and 3C). Overall, this data suggests that mesoderm internalization promotes the synchrony of MD1 divisions.

**Figure 3.**
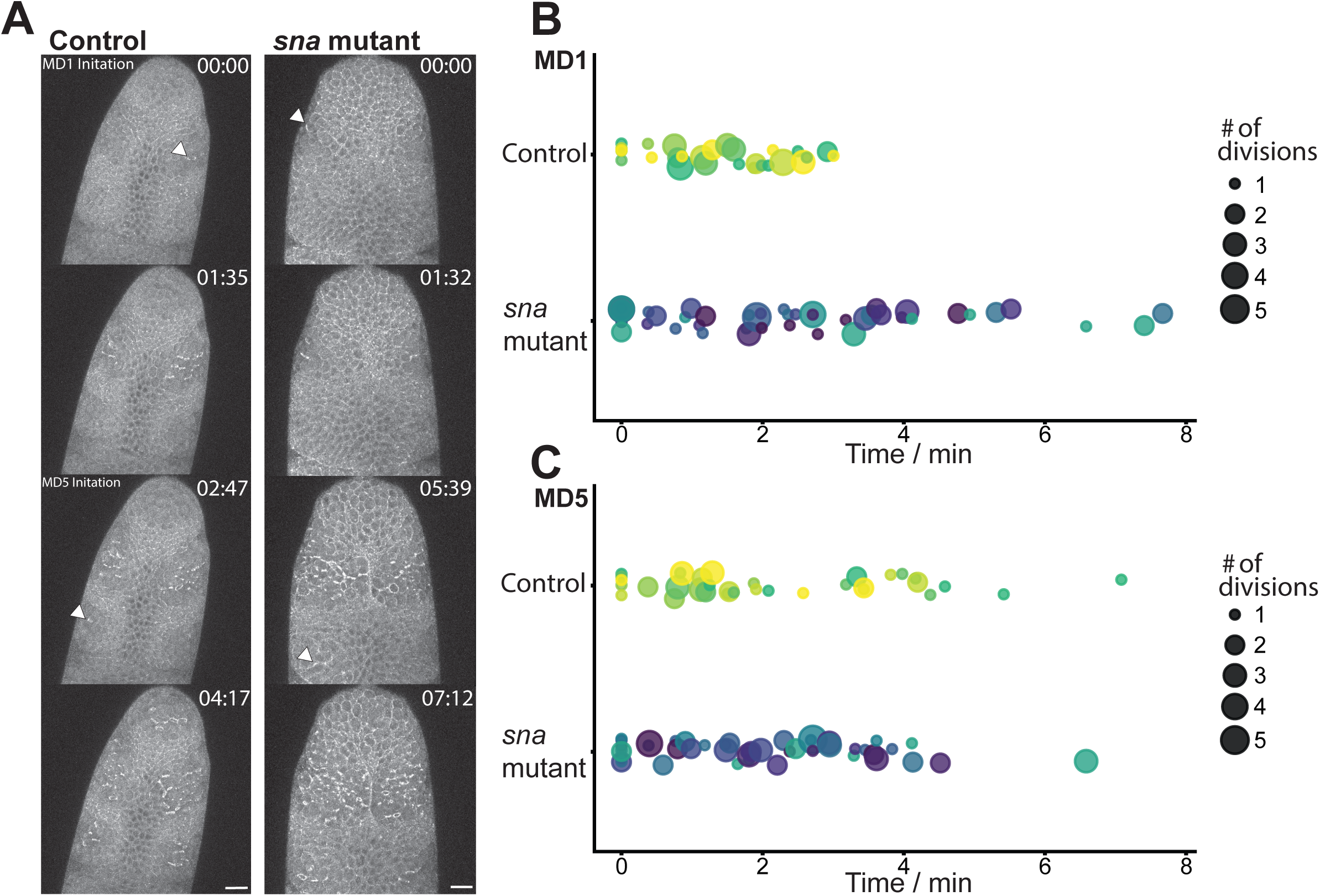
*snail* mutant disrupts mitotic synchrony of MD1. (A) Montage of time-lapse images of embryos expressing Squ::GFP. Compared are control Squ::GFP and snail mutant embryos. Scale bar = 25µm. (B) Synchrony of MD1 was driupted in the first 10 divisoins in the right side of 8 snail mutant embryos compared to 5 control. Snail mutants showed a 109% increase in division variability (p=0.016) and a difference in mean division time of +1.02 mins, nearing significance (p=0.068). Whereas (C) MD5 divisons were unaffected. Variabilty: (p=0.9720). Mean division time (p=0.721).

## Discussion

Previous research has shown that mitotic entry during gastrulation is closely associated with *cdc25^string^* expression, a phosphatase that regulates Cdk1 activity (Edgar and O’Farrell, 1990; Di Talia and Wieschaus, 2012). Our study shows that the mechanical forces generated during gastrulation in *Drosophila melanogaster* contribute to the timing of cell division in mitotic domains. We showed that forces generated during gastrulation influence division timing by: 1) disrupting the adherens junctions, which we also showed affects the timing of cellularization, 2) disrupting mesoderm invagination in a *sna* mutant, which disrupted the synchrony of mitotic entry in MD1, but not MD5.

### Adherens junction disruption has global effects on morphogenetic timing

Adherens junctions connect the neighboring cells physically, which promotes force transmission across epithelial tissues (Cox *et al*., 1996; Tepass *et al*., 1996). Disrupting these junctions not only affects physical connectivity but also impairs the cell’s ability to sense and respond to their mechanical environment (Lecuit and Yap, 2015). When disrupting the adherens junction component, α-Catenin, with RNAi we observed that cellularization proceeded at faster rate when compared to the control, suggesting that early morphogenetic processes may be sensitive to physical junctional integrity even before blastoderm formation. There are non-junctional E-cadherin structures present in the syncytial membrane, and it is possible that this affects the availability of membrane for cellularization furrow extension and/or the compartmentalization of the furrow canal (Sokac and Wieschaus, 2008; Figard *et al*., 2013). Future work is needed to determine how α-Catenin regulates cellularization and whether MD division timing is connected to the end of cellularization.

Beyond these early effects, disruption of α-Catenin also alters tissue architecture and cell division dynamics during gastrulation. We observe abnormal tissue architecture such as extra folds in the posterior region. Furthermore, we show that α-Catenin RNAi slows the progression of the first 10 divisions in MD1. These phenotypes are consistent with the loss of physical coupling in the tissue and suggest that forces are involved in coordinating cell division with other morphogenetic events during gastrulation. To test this, we investigated whether blocking mesoderm internalization similarly delayed the progression of MD divisions.

### The *snail* mutant disrupts mitotic entry timing in dorsal head

During mesoderm invagination, tissue movement and cell shape changes generate mechanical forces that get transmitted globally in the embryo. These mechanically generated forces have been studied extensively in the field, showing apical constriction that changes tissue curvature leading to the gastrulation in *Drosophila* (Martin, 2020). Similarly, previous studies have shown that these forces are not only restricted to the ventral furrow but also contribute to the global tissue movement, where changes in myosin pools play a major role in flow field configuration (Streichan *et al*., 2018). However, the role of these forces in regulating cell division has not been well studied. Our finding suggests that forces generated during mesoderm invagination plays major role in mitotic division in neighboring tissues. These mechanisms of physical connectivity allow cell to divide in a spatially and temporally regulated manner. Past studies have shown that *cdc25* regulates the timing and location of cell division in *Drosophila* embryo (Edgar and O’Farrell, 1989). In addition to this, our result shows that mechanical cues function alongside genetic program that regulate the timing of cell division. Our result where we show that disrupting the force transmission impacts the timing of cell division implies that patterning during development alone does not regulate the spatial and temporal resolution of cell division.

Our next result from *snail* mutant affecting the mesoderm invagination but not cell-cell junction, shows that inhibiting mesoderm invagination disrupts the progression of cell division within mitotic domains. Because *snail* is expressed exclusively in the mesoderm at this stage, this suggests that mechanical forces generated during gastrulation non-autonomously regulate cell division timing. Recent study has also shown that there is extensive coordination between cell shape changes, tissue rearrangement, and cell division during morphogenesis, claiming that forces are central player when viewing epithelium within the continuum mechanics framework (Stern *et al*., 2022). Although the *snail* mutant addresses the issue of cell-cell connectivity and only disrupt the mesoderm invagination, *snail* is a transcription factor that regulates many other genes that are involved in gastrulation (Leptin, 1991; Ip *et al*., 1992). Therefore, the changes in the timing and progression of cell division seeing in *snail* mutant might not be solely due to the loss of mechanical forces generation, rather it could be due to the alteration of another developmental program.

### Limitations of study

Our findings shows that the role of mechanical forces generated during gastrulation and cell-cell junction are important factors in regulating the timing of cell division. The mechanism through which mechanical forces regulate cell division remains an open question. It is possible that mechanical forces regulate the activity of mechanosensitive signaling pathways that regulate the G2/M transition. Although our results demonstrate the relationship between mechanical forces and cell division, the molecular pathways through which this regulation occurs remain unclear. Further studies involving force measurement and mechanosensitive reporters are necessary to elucidate the downstream pathways of forces generated during gastrulation in regulating the cell division.

In α-Catenin RNAi embryos, differences in knockdown efficiency between embryos lead to variation in phenotype severity. While the overall trend in the timing of cell division supports the central role of adherens junctions and mechanical forces in the regulation of cell division timing, it is possible that differences in timing of cell division result from embryos that exhibit a stronger knockdown phenotype. Additionally, our experiments did not measure the forces directly in the embryo, rather force generation and transmission were inferred through genetic perturbation. Further studies, quantifying forces can answer questions such as how much force is required for cells to enter into mitosis, revealing the casual relationship between the two. Additionally, disrupting the adherens junction through the depletion of α-Catenin, also disrupts cell adhesion, epithelial integrity, mechanosensory signaling, and cytoskeletal organization. Therefore, the changes in the timing of cell division that we observed may not be solely due to impaired force transmission.

## Materials and Methods

### Fly stocks and genetics

All *Drosophila melanogaster* stocks were maintained at 25 °C. To generate embryos for analyzing the effects of the *snail* mutation, a two-generation genetic cross strategy was employed. At first, a virgin flies carrying a Histone::GFP on the third chromosome was crossed to a mutant male flies carrying *halo snail* mutations balanced over the CyO::Sqh-GFP balancer on the second chromosome (*halo snail/CyO::Sqh-GFP*) (Martin *et al*., 2009). From the F1 generation embryos, halo mutant embryos were selected and imaged to ensure that we selected homozygous *halo sna/halo sna* mutant embryos.

To investigate the effects of α-Catenin depletion during early embryogenesis, a two-generation cross strategy was employed. At first, virgin females carrying α-Catenin RNAi (TRiP.HMS00317) balanced over TM3 on the third chromosome (*α-Catenin RNAi/TM3*) were crossed with male flies of the genotype *mat67,sqh-mCherry/CyO; Histone::GFP/TM3* (Ni *et al*., 2011). From the F1 generation, virgin females were selected based on the absence of both balancer chromosomes (CyO and TM3) to isolate individuals of the genotype *+/mat67,sqh-mCherry; α-Catenin RNAi/Histone::GFP*. F1 virgin females were then crossed to males homozygous flies for Histone::GFP on the third chromosome to generate F2 maternally depleted embryos.

### Live imaging (Confocal and Brightfield Microscope)

For live imaging, embryos were collected on apple juice agar plates and aged to the desired developmental stage at 25 °C incubator. Embryos were dechorionated by immersing them in 50% household bleach for 2 minutes. Following dechorionation, embryos were thoroughly rinsed with distilled water to remove residual bleach.

Dechorionated embryos were transferred onto a glass slide previously coated with embryo glue. The glue was prepared by dissolving double-sided adhesive tape in heptane and applying it evenly to the slide surface. Embryos were aligned ventral side down using a fine paintbrush under a light microscope. To create an imaging chamber, two No. 1.5 coverslips pre-coated with the same embryo glue were placed on either side of the embryo row to serve as spacers. A No. 1 coverslip was then gently placed on top, bridging the spacer coverslips and enclosing the embryos. The chamber was sealed and filled with Halocarbon 27 oil to maintain embryo hydration. Fluorescently labeled embryos were imaged using a Zeiss LSM 710 confocal microscope. Appropriate lasers were used to visualize specific fluorophore. All imaging was carried out at room temperature (around 23-25 °C).

For the brightfield imaging, embryos were placed onto a glass slide with Halocarbon 27 oil and imaged them under brightfield optics using a Nikon 10x and 20x numerical aperture objective. Embryos were imaged at room temperature (23–25 °C) using a time-lapse setup on a brightfield microscope, and images were acquired at regular intervals of 2 minutes throughout the process.

### Image processing and statistical analysis

All image processing were performed using Fiji. Brightness and contrast adjustments were applied uniformly across entire images to enhance visualization of nuclear or Myosin II signals with identical adjustments applied to both control and experimental conditions to ensure consistency and comparability. Images used in figures represent maximum intensity projections or single z-slices, as indicated in figure legends. All quantitative analyses were conducted on unprocessed raw images.

All quantitative measurements and statistical analyses were performed using GraphPad Prism. Statistical significance was assessed using appropriate tests (Mann– Whitney test and ANOVA) depending on data distribution and experimental design. A p-value of less than 0.05 was considered statistically significant. Details of the statistical tests used, sample sizes, and error bars are indicated in the corresponding figure legends.

### Cell division and cellularization timing analysis

To analyze the timing of cell divisions, embryos expressing Histone::GFP were used to visualize nuclear dynamics. Embryos were selected around one hour after egg laying, prior to the onset of the 13th nuclear division cycle. The onset of cycle 13 in the embryo was identified by the observation of the first anaphase in the field of view, which served as a temporal reference point and called as the 13th anaphase.

For the analysis of Cephalic Furrow (CF) formation, the initiation of furrow invagination was well-defined as the time point when the first nucleus disappeared from the field of view due to internalization at the presumptive CF site.

The timing of anaphase onset in Mitotic Domains (MDs) 1 and 5 was determined by observing the first clear anaphase event in each domain within the field of view. The timing of the first 10 cell divisions was recorded by manually scoring the first 10 nuclei to enter anaphase on one side of the embryo within MD1 and MD5.

To measure the timing of cellularization, the start of cellularization was defined as the time point at which the initial inward movement of the furrow canal in the brightfield images. The end of cellularization was marked by the time at which the invaginating membranes reached the yolk sac, visibly completing the enclosure of nuclei into individual cells. The timing of cellularization was quantified manually by analyzing the frames corresponding to the start and the end of cellularization as described earlier.

## Notes

### Competing Interest Statement

The authors have declared no competing interest.

### Funding

- National Institutes of Health: R35 GM144115

